# Pluripotent Stem Cells Can Be Isolated from Human Peripheral Nerves after *in vitro* BMP-2 Stimulation

**DOI:** 10.1101/598433

**Authors:** Ren-Yi Sun, Michael H. Heggeness, Tanghong Jia, Sunaina Shrestha, Bradley Dart, Shang-You Yang

## Abstract

We have recently identified a population of cells within the peripheral nerves of adult mice that can respond to BMP-2 exposure or physical injury to rapidly proliferate. More importantly, these cells exhibited embryonic differentiation potentials that could be induced into osteoblastic and endothelial cells in vitro. The current study examined human nerve specimens to compare and characterize the cells after BMP-2 stimulation. Fresh pieces of human nerve tissue were minced and treated with either BMP-2 (750ng/ml) or vehicle for 12 hours at 37°C, before digested in 0.2% collagenase and 0.05% trypsin-EDTA. Isolated cells were cultured in restrictive stem cell medium. Significantly more cells were obtained from the nerve pieces with BMP-2 treatment in comparison with the non-treated controls. Cell colonies were starting to form at day 3. Expressions of the 4 transcription factors Klf4, c-Myc, Sox2 and Oct4 were confirmed at both transcriptional and translational levels. The cells can be maintained in the stem cell culture medium for at least 6 weeks without changing morphologies. When the cells were switched to fibroblast growth medium, dispersed spindle-shaped cells were noted and became fibroblast activated protein-α (FAP) positive following immunocytochemistry staining. The data suggested that human peripheral nerve tissue also contain a population of cells that can respond to BMP-2 and express all four transcription factors KLF4, Sox2, cMyc, and Oct4. These cells are capable to differentiate into FAP-positive fibroblasts. It is proposed that these cells are possibly at the core of a previously unknown natural mechanism for healing injury.

## Introduction

The potential for stem cells to treat human disease is rightly perceived to be vast. Embryonic stem cells (ESCs) from inner cell mass of mammalian blastocyst that have unlimited self-renewal and pluripotency are capable of differentiate into ectodermal, mesodermal, and endodermal cells [1, 2]. Based on the stud of previous, the ESCs behave undifferentiated morphology [3]. Although there are numerous ongoing studies to investigate the therapeutic potentials of human embryonic stem cells (hESCs) for type I diabetes (T1D), heart failure, Parkinson’s disease and inherited or acquired retinal degenerations [4], challenges remain to be conquered in clinical development of hESCs such as legal and ethical issues, immune rejections, and differentiation difficulties [3]. Somatic cells can be introduced to transform into a state of pluripotency [3]. The brilliant work of Drs. Yamanaka and Takayashi [5, 6] demonstrated that pluripotent cells can be created from adult differentiated cells by the virally induced manipulation of nuclear genes to force expression of 4 specific transcription factors, octamer-binding transcription factor 4 (OCT4), sex determining region Y-box 2 (SOX2), Krüppel-like family of transcription factor 4 (KLF4) and c-Myc that will convert the cells to pluripotency. Unfortunately, this process creates a very real risk of malignant transformation, and does not solve the issue of immune rejection, as the cells are “non-self”. Another drawback is that such cells are expensive to create and assessing the cells for risk of malignant transformation adds further time and expense. At best it can diminish this risk, but does not eliminate it [7, 8]. We have serendipitously discovered a population of pluripotent cells that reside in a quiescent state within mouse peripheral nerves [9, 10]. When the nerves are stimulated with physical insult (including mechanical compression or stretching, exposure to blood, and electrical or cytokine BMP2 stimulation) a massive proliferation of cells within the nerve results with a rapid egress of the cells into the surrounding tissues. These proliferating cells uniformly exhibit expression of the 4 critical genes associated with pluripotency; Sox2, Oct4, c-Myc, and Klf4 as demonstrated by double stain immunohistochemistry and by Real Time Polymerase Chain Reaction (RT-PCR) with appropriate primers [10]. They are readily cultured in restrictive media, adhere to substrate, and appear motile. They have been successfully differentiated into cells of the three primary germ layers: mesoderm (osteoblasts and endothelial cells) [9], endoderm (Definitive Endoderm), and ectoderm (Primitive Nerve Cells) in rodents (unpublished data). We have termed these cells NErve Derived Adult Pluripotent Stem cells, or NEDAPS cells.

Indeed, there are too many instances that the biological data from rodent experiments did not conform to human tissue responses. The objective of this study was to examine the characteristics and differentiation potential of these pluripotent stem cells, remarkably, obtainable from adult human peripheral nerves.

## Materials and Methods

### Human peripheral nerve tissue treatment

This study was exempted by the institutional Review Board (IRB) as a non-human investigation. Fresh human peripheral nerves were obtained as surgical waste in a completely anonymous fashion from 3 serious trauma patients during limb amputation. This was tissue that would normally have been discarded. No identifying features of the specimens was shared or recorded. The peripheral nerve tissues were stored in sterile normal saline on ice and transferred to the research lab within one hour from the amputated limb. The nerves were immediately minced and put into a 6-well culture plates (CytoOne, USA Scientific, Ocala, FL, USA) with DMEM medium and 275 ng/ml of rhBMP-2 (InFuse™, Medtronic, Memphis, TN) was added to each well. The minced nerve tissue was incubated overnight at 37 °C before cell isolation.

### Mouse sciatic nerve treatment

Thirty (30) BALB/c mice at age of 10 weeks were used for this study. The institutional Animal Care and Use Committee (IACUC) approved all the animal protocols, and the surgical procedures were performed as reported previously {Heggeness, 2017 #50}. Briefly, both hind legs of mouse were shaved and disinfected after anesthesia by intraperitoneal injection of 8mg/kg Xylazine and 90mg/kg Katamin. Under strict sterile condition, sciatic nerve was surgically exposed through lateral incision, and 20 µl of rhBMP-2 applied to the nerve. The wound was closed in layers, and the animals were kept for 20 hours before sacrifice for the sciatic nerves harvest.

### Cell isolation and Culture

Proliferating NEDAPS cells were isolated the same as reported previously for mouse NEDAPS cells [10]. Briefly, the minced nerve pieces were pelleted by centrifugation at 500xg for 5 minutes and followed by digestion in 0.2% (0.27 U/ml) collagenase (Worthington Biochemical Corp) at 37°C for 90 minutes. Equal volume of 0.05% trypsin-EDTA solution was then added for 5 minutes with agitation. The enzyme digestion was stopped by addition of heat-deactivated fetal bovine serum (FBS) and the mixture was filtered through a 100μm-sized mesh before centrifugation at 500xg for 10 min to pellet cells. The cells were seeded into 6-well culture dishes, or 8-well chamber-slides in the restriction stem cell medium with 20% Knockout serum replacement (KSR, Gibco), 100μM MEM non-essential amino-acid solution (Gibco), 1x GlutaMAX™-I (Gibco); 55μM β-mercaptoethanol (Gibco), and 20 ng/ml human leukemia inhibitory factor (LIF, Gibco). The cells were cultured at 37°C, 5%CO_2_ atmosphere. For fibroblast induction, the human NEDAPS cells were switched to the fibroblast induction medium that contains 20 ng/mL recombinant human FGF-2 (rhFGF-2), 50 µg/mL Ascorbic Acid, and 10% FBS in DMEM.

### Real-time PCR (RT-PCR) test

For RNA isolation, the cells were resuspended and lysed in TRIzol solution (Life Technologies, Carlsbad, CA, USA), and then went through chloroform separation and isopropanol precipitation following the protocol described by Chomczynski [11, 12]. Complementary DNA (cDNA) was made by reverse transcription in the 40 μl mixture as follows: 4 μl 10 × PCR Rxn buffer (-MgCl_2_) (Invitrogen, Grand Island, NY, USA), 4.4 μl MgCl_2_ in the concentration of 5.5mM (Invitrogen), 8 μl deoxynucleoside triphosphates in the concentration of 500 μM (Invitrogen), 2 μl RNase inhibitor in the concentration of 0.5 U/μl (Invitrogen), 2 μl random hexamers in the concentration of 2.5 μM (Invitrogen), 0.25 μl reverse transcriptase in the concentration of 1.25 U/μl (Invitrogen), 9.35 μl DNAse, RNAse free water (Invitrogen) and 0.5 μg of extracted RNA. The reactions were in fast reaction tubes on a Veriti 96-well Thermal Cycler (Applied Biosystems, Foster City, CA, USA). The temperature cycle was: at 25°C for 10 minutes, 48 °C for 30 minutes and 95 °C for 5 minutes. RT-PCR was run on a StepOnePlus RT-PCR System (Applied Biosystems) for forty cycles in fast optical 96-well reaction plate. The 25 μl reaction mixture was mixed by 12.5 μl SYBR Green PCR Master Mix (Applied Biosystems), 6 μl cDNA, 5.5 μl DNAse, RNAse free water (Invitrogen) and 400 nM tested gene primer pairs. The RT-PCR System can automatically record all the fluorescent signals dynamically. We employed Primer3 program (http://bioinfo.ut.ee/primer3/) to design the primer pairs for each target gene[13, 14] and made by Sigma-Genosys (Woodlands, TX, USA). The gene sequences are described as in Table 1.

**Table 1.**
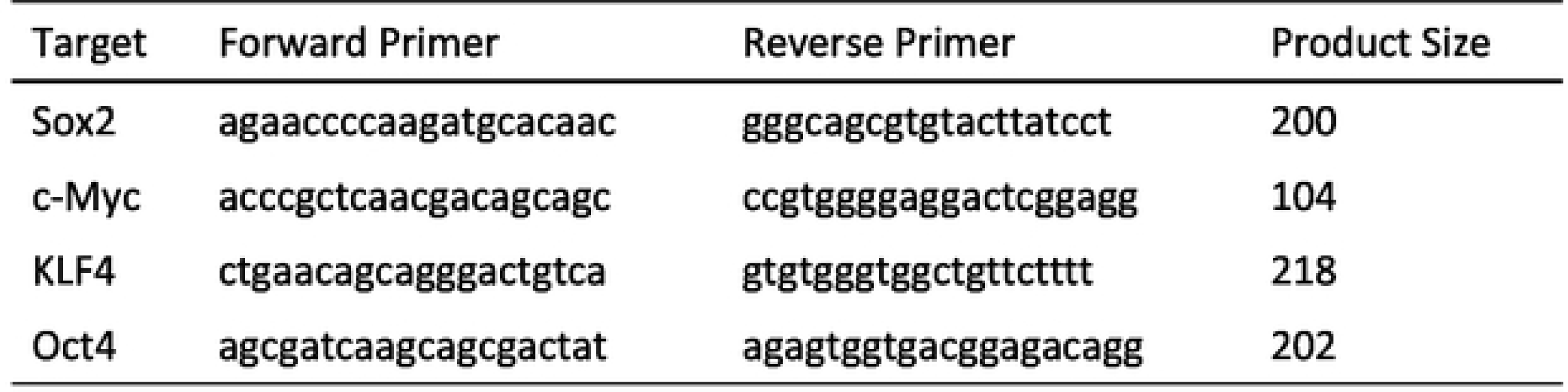
Primers Utilized for RT-PCR Amplification of human specimens

### Immunocytochemistry staining and image acquisition

The immunocytochemistry staining technique for the four stem cell markers (c-Myc, Sox2, KLF4, Oct4) has been reported previously [9, 10]. The following primary antibodies were used as pairs for double staining: goat anti-Sox2 (Santa Cruz), goat anti-KLF4 (R&D), rabbit anti-c-Myc (Abcam), goat and rabbit anti-Oct4 (Abcam), rabbit anti-Myelin (Abcam). Cells were double stained with goat and rabbit first antibody pairs (Sox2 + c-Myc, KLF4 + c-Myc, Sox2 + Oct4, KLF4 + Oct4, KLF4 + Myelin, Oct4 + Myelin). For fibroblast differentiation, rabbit anti-FGF-2 (basic fibroblast growth factor, Abcam) and rabbit anti-mouse FAB (fibroblast activation protein, Abcam) were used. Following washes, the diluted secondary antibody (1:200) in 1 × PBS with 1% BSA was added onto the cells for 1h at room temperature in the dark. The secondary antibodies we utilized were donkey anti-rabbit IgG conjugated with Alexa Fluor 594 (Life Technologies) and donkey anti-goat IgG with Alexa 488. DAPI Fluoromount G (Southern Biotech, Birmingham, AL, USA) was used to mount coverslip and to counterstain cell nuclei. Fluorescent images of the cells were acquired under a Nikon E800 fluorescence microscope (Nikon, Japan), by a Coolsnap EZ CCD Camera (Photometrics, Tucson, AZ, USA) and analyzed using a MetaMorph image analysis software (Molecular Devices, San Francisco, CA, USA). Variously stained images were overlaid by the image analysis software to illustrate the co-localization patterns.

### Statistical analysis

The data of comparative gene expression of the 4 stem cell transcription factors and the fibroblast markers between groups and immunocytochemical positive cells were all recorded, and expressed as mean ± standard deviation. The data among the groups were analyzed by one-way analysis of variance (ANOVA) followed by Bonferroni post-hoc test (equal variances assumed). Statistical probability of P < 0.05 was considered as statistical significance. The software for statistical analysis was IBM SPSS Statistics (Armonk, NY, USA).

## Results

### Morphology of the human NEDAPS cells

Similar to the mouse NEDAPS cells, the human cells readily adhered to the culture plates or chamber-slides and maintained a polygon-shaped morphology (Figure 1A). They also did not require feeder cells to survive. However, in contrast to the rodent cells [10] which were remaining dispersed on the culture plate, the human cells tended to gather to form colonies. It appears that the motile cells formed clusters within 24 to 48 hours in the stem cell medium, and stayed in colonies during proliferation. Indeed, the cells in colonies continued to divide and could reach several cell layers (Figure 1B).

**Figure 1:**
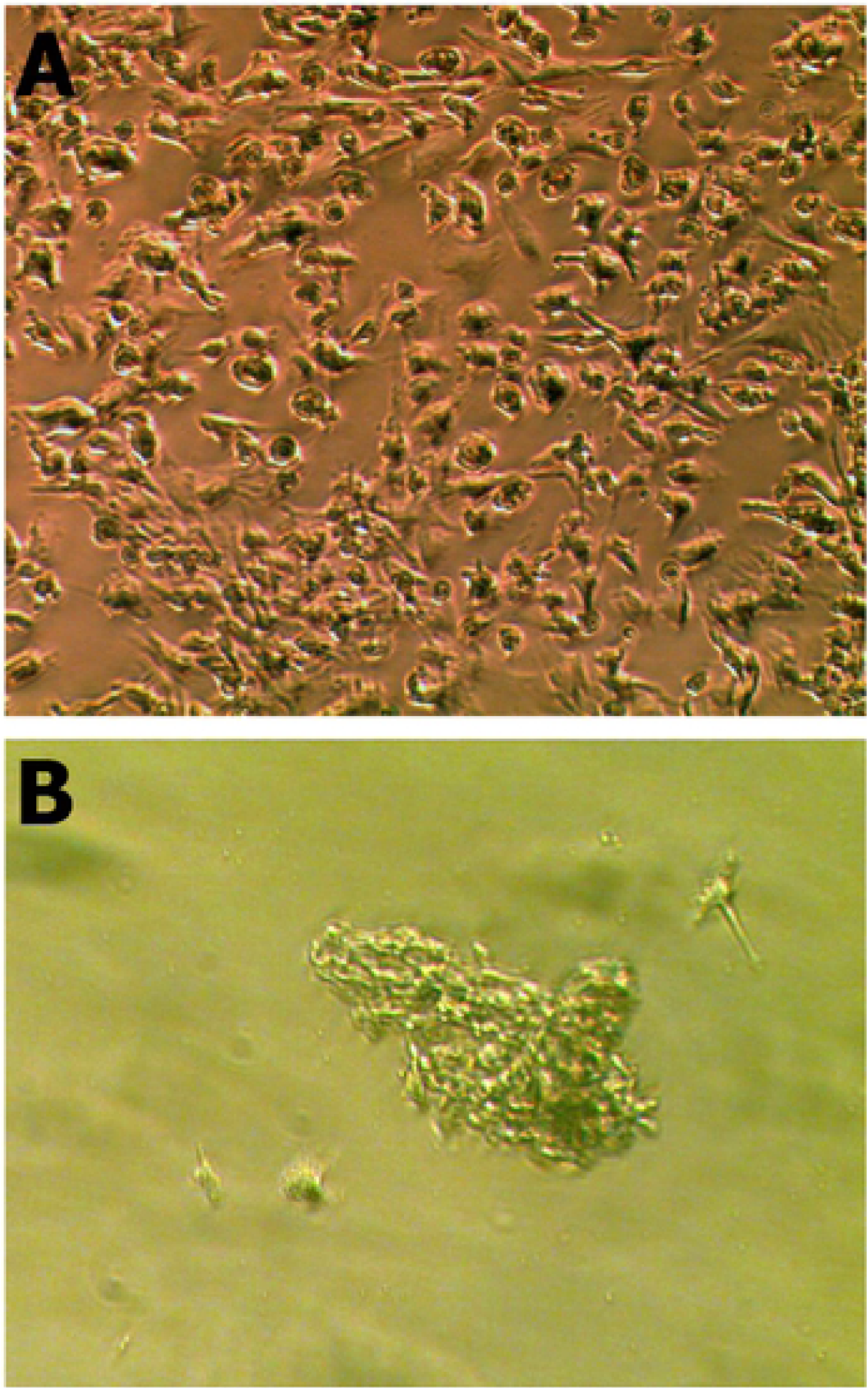
NEDAPS cells isolated from human peripheral nerve tissue and placed in cell culture dish with complete stem cell restrict medium for 24 hours (A), and 7 days (B).

### Expression of stem cell markers

Immunofluorescent microscopy clearly showed that over 90% of the isolated human nerve cells were simultaneously expressing the four transcription factors Sox2, Oct4, c-Myc and Klf4 after 5 days in culture in the restrictive stem cell medium. It is also noticeable that the double staining illustrates the co-expressions of the stem cell markers in various pairs (Figure 2).

**Figure 2:**
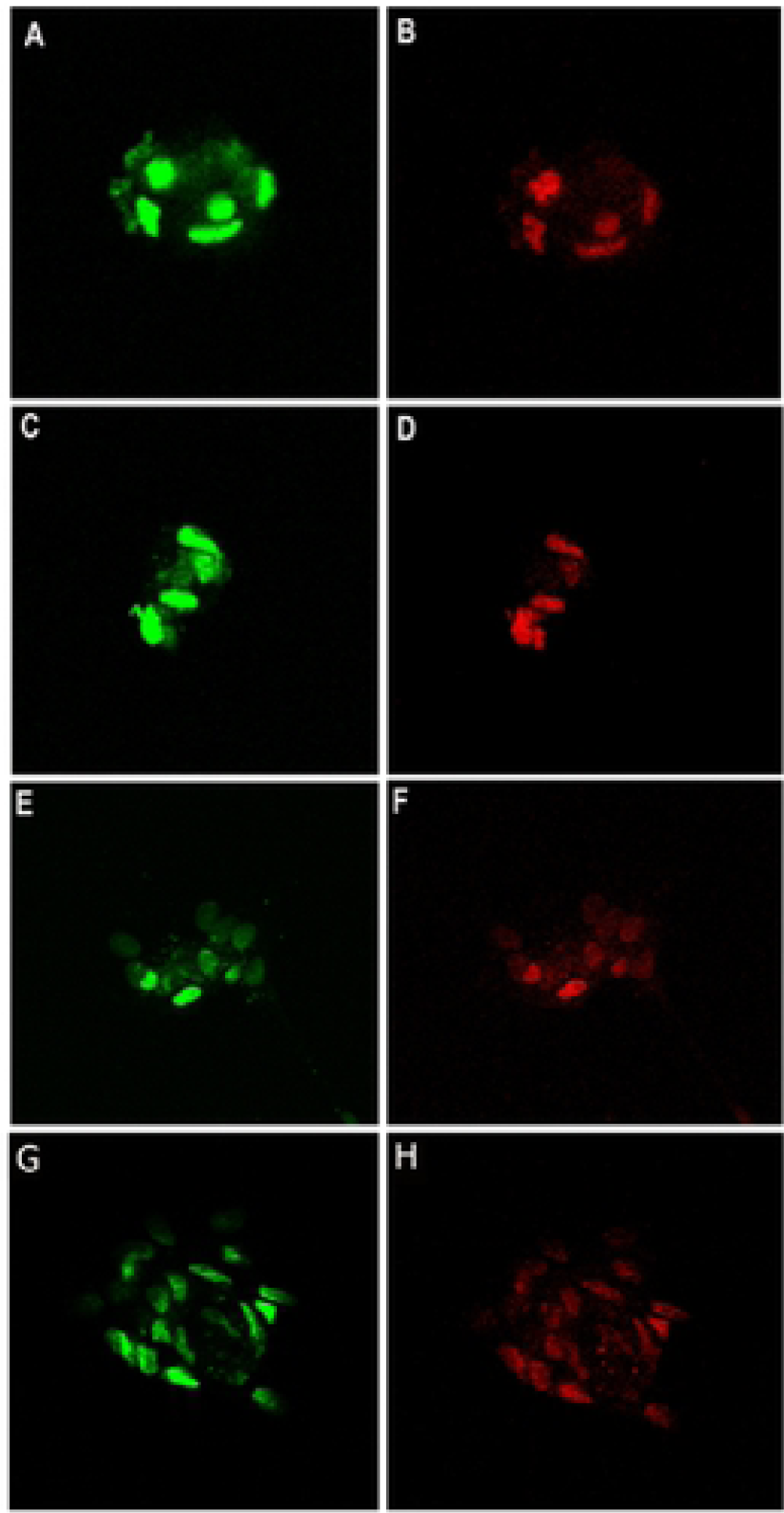
lmmunofluorescent staining was performed to identify the pluripotent stem cell markers, and visualized under a confocal microscope. Cells were double stained with a pair of primary antibodies raised in different species, and probed with Alexa Fluor^®^488 secondary Ab (green) and Alexa 594 2nd Ab (red): (A and B) Sox-2 and Klf-4; (C and D) Oct-4 and c-myc; (E and F) Sox-2 and c-myc; (G and F) Klf-4 and Oct-4.

Real time PCR was performed to examine the mRNA expressions of Sox2, Oct4, c-Myc, and Klf4 of these cells. It is convincing that strong expressions of all the 4 transcription factors in the human nerve specimens treated with BMP-2, significantly more than those obtained from non-BMP2 treated nerve specimens (p<0.05). Some of the samples following real-time PCR were electrophoresed on an agarose gel to reveal the correct-sized PCR product of the transcription factors (Figure 3).

**Figure 3:**
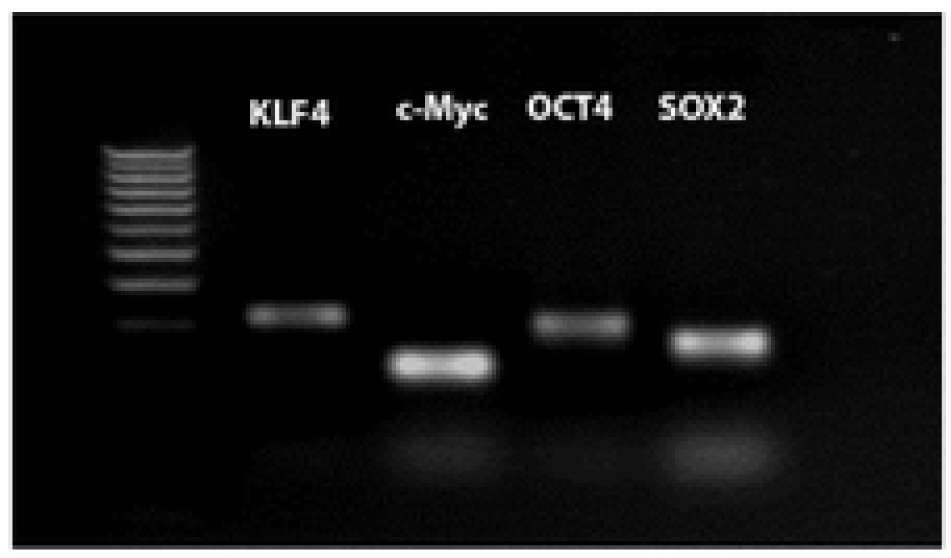
Electrophoresis of PCR product after the real-time PCR with specific primers for the 4 transcription factors, on the BMP-2 treated cells isolated from human peripheral nerves. The first lane shows a 100pb DNA ladder.

### Differentiation to fibroblasts

When the human NEDAPS cells were changed to the complete fibroblast induction medium, they rapidly dispersed from their small colonies and assumed the classic “spindle shape” of motile fibroblasts within 24 – 48 hours (Figure 4). Immunohistochemical staining against fibroblast activation protein (FAP) confirmed that they become FAP-positive.

**Figure 4:**
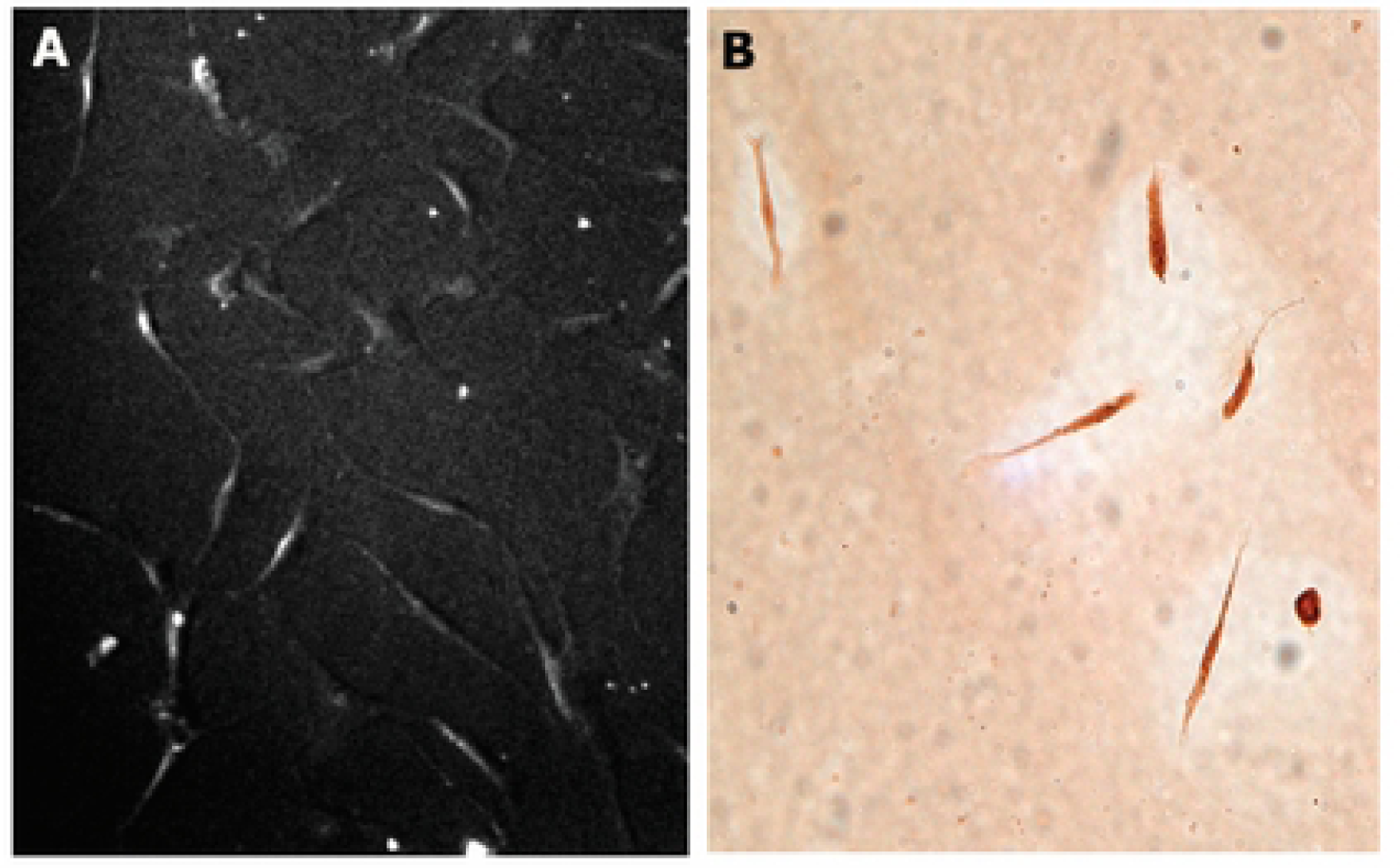
Human NEDAPS cells were morphologically changed to spindle-shaped fibroblastic cells (A), and stained strongly positive for fibroblast activation protein alpha positive (B).

## Discussion

Bone morphogenetic proteins (BMPs), first described by Marshall Urist in 1965 [15], are able to stimulate the stem cell to differentiate into osteoblast cell, and are considered to be the most important growth factors in bone formation and fracture healing. As members of the TGF-β superfamily, the details of their status as intercellular signal transduction are still under study. Among the more than 20 different human BMPs identified until now, only the BMP-2 and BMP-7 are approved by the Food and Drug Administration (FDA) for clinical use. In recent years, more and more Orthopedics surgeons are likely to apply BMP-2 to a variety of different clinical applications for its osetoinductive function [16]. With the increase in clinical use of BMP-2, many associated adverse events have been reported, most of them related to the use of BMP2 (Infuse) in proximity to nerves [17]. We speculate that the use of BMP2 in proximity to nerves at vastly supraphysiologic levels will likely lead to nerve dysfunction as the supraphysiologic production of stem cells will result in a likelihood of permanent nerve dysfunction [10].

In the effort of using mouse model to investigate potential complications of BMP-2 to peripheral nerves during bone healing applications, an intriguing finding was noticed that a population of cells was proliferated within the nerve. Indeed, these cells were able to express the 4 transcription factors c-Myc, Klf4, Sox-2, and Oct-4 [10], suggesting their pluripotent potentials. Further investigation also confirmed that physical injury to peripheral nerves such as compression also induced the same phenomena of cell proliferations [10]. In the current study, it is very interesting to note that the same population of cells can be isolated from human peripheral nerve tissue after in vitro incubation with small amount of BMP-2, although the cell growth pattern of human nerve induced cells appear different compare to the mouse NEDAPS cells, the human tibial induced cells all huddled together and grew concentric but the mouse NEDAPS cells were all dispersive.

The immunocytochemistry and RT-PCR both verified new gene expression (Sox2, KLF4, c-Myc, Oct4), and these four transcription factors were regarded as exclusive for pluripotent stem cells. Observed utilizing fluorescence microscope, the slides which were treated by immunochemical technology clearly displayed the four stem cells’ exclusive marker. Real-time RT-PCR technique conveniently confirmed expressions of the four target genes (Sox2, KLF4, c-Myc, Oct4) and the agarose electrophoresis revealed this result.

To illustrate the differentiation potential of the human NEDAPS cells, we switched the cell culture medium to fibroblast growth medium containing recombinant human fibroblast growth factor-2 (rhFGF-2). The cell morphology was quickly changed with 2-3 days, and IHC staining confirmed expression of fibroblast activation protein (FAP-α). FAP-α has been identified as a fibroblast marker that is over-expressed in activated fibroblasts or mesenchymal stromal cells [18], the differentiation ability of the human nerve-derived NEDAPS cells will be further investigated to explore the clinical significance.

The stem cells, which are also immature cells with self-renewal capacity, are able to differentiate into many kinds of somatic cells depending where they come from [19]. However, if so used as an allograft cell-based therapy, the recipients would need to be under an immune-suppressive therapy in order to prevent immune rejection [7, 8].

We are excited to report on this very recently discovered source of pluripotent cells which would seem to have an exciting potential for human cell-based therapies. These cells are notably different from embryonic stem cells or the induced pluripotent stem cells (iPCs), with remarkably distinct appearance and behaviors compared to previously described “true embryonic stem cells”. Importantly, these peripheral nerve-derived cells seem to have the natural function of accomplishing tissue repair and their “niche” within peripheral nerves provides a welcome insight into the pathophysiology of the wound healing problems associated with peripheral neuropathies including leprosy, Diabetes mellitus and tertiary syphilis. We speculate that the loss of healthy nerves in these patients means that the NEDAPS cells are no longer available to heal such wounds. This insight may open the door to all manner of cell-based treatments using self-to-self autografts of pluripotent NEDAPS cells or differentiated cells therefrom. This could potentially allow the harvest of a non-essential peripheral nerve such as a branch of the purely sensory sural nerve, of the anterior intraosseous nerve (or many others) to perhaps have a very wide application for tissue regenerations.

In conclusion, we suggested in this study that the human peripheral nerves also contain normally quiescent NEDAPS cells. Current investigation is focused on further documenting their differentiation potentials and to exploring their therapeutic potential in bone and other tissue repair and regeneration models. We suggest that these cells are possibly at the core of a previously unknown natural mechanism for healing injury.

## Acknowledgments

The authors wish to acknowledge valuable technical assistance of Ms. Zheng Song in the early stage of the investigation.

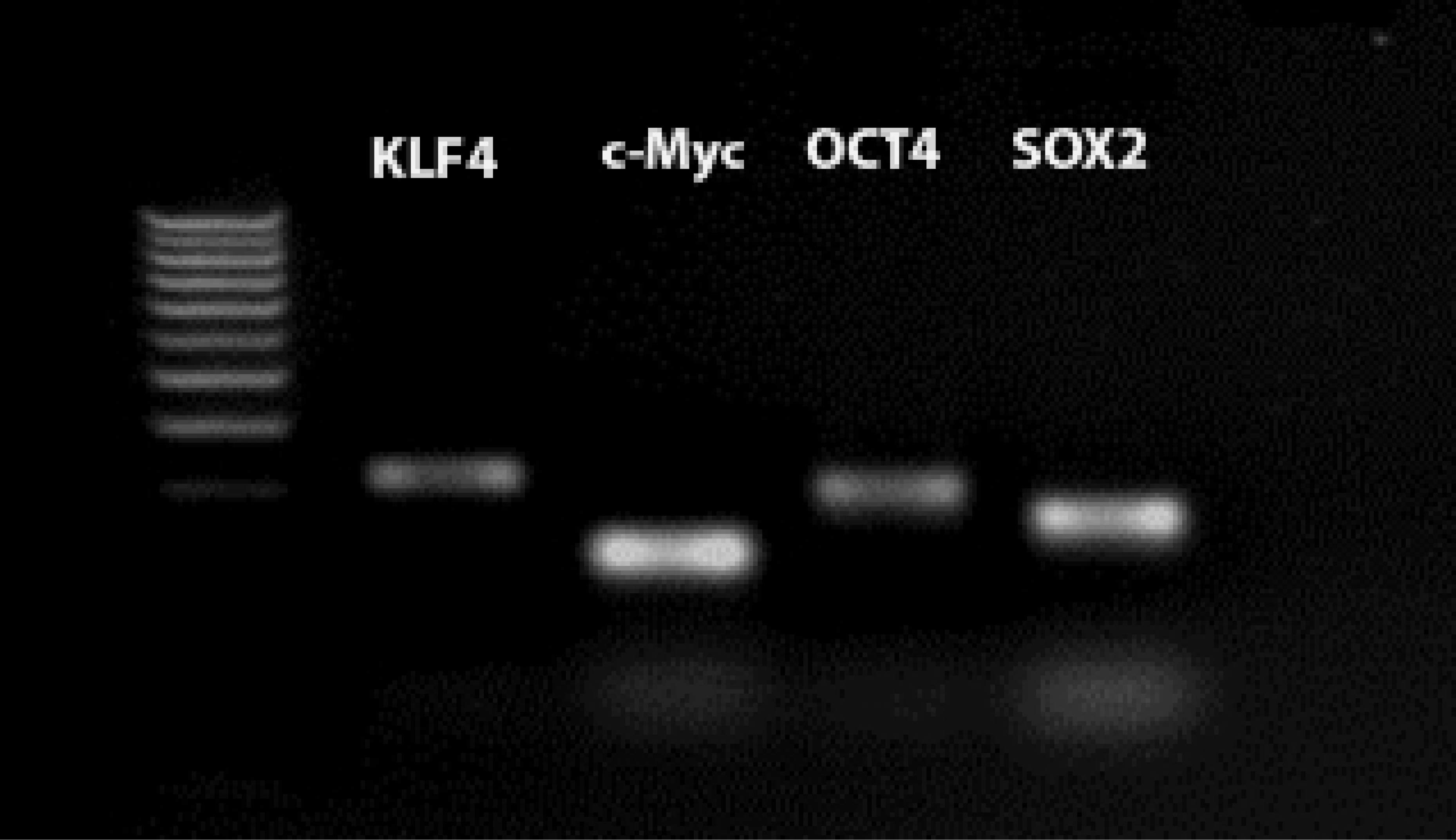

